# Evaluating treatments for the protection of grapevine pruning wounds from natural infection by trunk disease fungi

**DOI:** 10.1101/2024.02.27.582328

**Authors:** C. Leal, R. Bujanda, B. López-Manzanares, S. Ojeda, M. Berbegal, A. Villa-Llop, L. G. Santesteban, J. Palacios, D. Gramaje

**Author notes:** Corresponding Author: David Gramaje.

## Abstract

Infection of grapevines by fungal pathogens causing grapevine trunk diseases (GTDs) primarily arises from annual pruning wounds made during the dormant season. While various studies have showcased the efficacy of products in shielding pruning wounds against GTDs infections, most of these investigations hinge on artificial pathogen inoculations, which may not faithfully mirror real field conditions. This study aimed to evaluate and compare the efficacy of various liquid formulation fungicides (pyraclostrobin + boscalid) and paste treatments, as well as biological control agents (BCA: *Trichoderma atroviride* SC1, *T. atroviride* I-1237, and *T. asperellum* ICC012 + *T. gamsii* ICC080), for their potential to prevent natural infection of grapevine pruning wounds by trunk disease fungi in two field trials located in Samaniego (Northern Spain) and Madiran (Southern France) over three growing seasons. Wound treatments were applied immediately after pruning in February. One year after pruning, canes were harvested from vines and brought to the laboratory for assessment of *Trichoderma* spp. and fungal trunk pathogens. More than 1,200 fungal isolates associated with five GTDs (esca, Botryophaeria, Diaporthe and Eutypa diebacks, and Cytospora canker) were collected from the two vineyards each growing season. Our findings reveal that none of the products under investigation exhibited complete effectiveness against all the GTDs. The efficacy of these products was particularly influenced by the specific year of study. A notable exception was observed with the biocontrol agent *T. atroviride* I-1237, which consistently demonstrated effectiveness against Botryosphaeria dieback infections throughout each year of the study, irrespective of the location. The remaining products exhibited efficacy in specific years or locations against particular diseases, with the physical barrier (paste) showing the least overall effectiveness. The recovery rates of *Trichoderma* spp. in treated plants were highly variable, ranging from 17% to 100%, with both strains of *T. atroviride* yielding the highest isolation rates. This study underscores the importance of customizing treatments for specific diseases, taking into account the influence of environmental factors for BCA applications.

## Introduction

Grapevine trunk diseases (GTDs) pose a significant threat to the sustainability of vineyards, serving as the primary cause of vine decline. These diseases, caused by various fungal pathogens, have a detrimental effect on plant productivity, ultimately resulting in long-term plant mortality (Gramaje et al. 2018). Consequently, a considerable proportion of vineyard acreage - sometimes exceeding 10% - needs replantation due to the loss of infected plants (Kotze et al. 2011; Bruez et al. 2013). Managing GTDs presents a challenge for winegrowers, nurseries, viticulturists, and scientists, given their complexity compared to other grapevine diseases like powdery and downy mildews (Bertsch et al. 2013). One intriguing and contentious aspect of GTDs lies in their undefined latency period often referred to as the asymptomatic phase (Hrycan et al. 2020). It is not uncommon for symptoms to manifest on one vine in a particular year while remaining absent in the following year, primarily due to environmental, climatic, and cultural factors (Sosnowski et al. 2011; Murolo et al. 2014). Consequently, accurately assessing the true incidence of GTDs in a vineyard during any given year becomes challenging, potentially leading to an underestimation of their overall impact.

There are various disorders classified as GTDs, each exhibiting distinct symptoms. Among the prominent GTDs affecting mature grapevines are Botryosphaeria dieback, Eutypa dieback, Phomopsis dieback, and esca disease (Gramaje et al. 2018). Additionally, recent studies have identified *Cytospora* spp. as a cause of dieback and wood cankers in grapevines (Lawrence et al. 2017). Research on these diseases has unveiled a significant correlation between their life cycles and specific cultural practices, particularly dormant pruning. It has been observed that winter pruning wounds act as the primary entry point for GTD pathogens in vineyards, primarily through the dispersal of airborne spores (Gramaje et al. 2018).

Fruiting bodies of GTD pathogens develop from dead/cankered wood, old pruning wounds, grapevine canes, crevices, cracks and on the bark of infected grapevines (Úrbez-Torres and Gubler 2011; Baloyi et al. 2016). The release of pathogen spores is predominantly influenced by rainfall and/or temperature events (Larignon and Dubos 2000; Eskalen and Gubler 2001; Úrbez-Torres et al. 2010; Van Niekerk et al. 2010; Valencia et al. 2015; González-Domínguez et al. 2021). Infection occurs when these spores land on exposed and vulnerable pruning wounds, germinate within the xylem vessels, and establish colonies within the vine spur, cordon, and trunk (Gramaje et al. 2018). The susceptibility of pruning wounds to GTD pathogens primarily depends on the timing of pruning and the period between pruning and potential infection occurrences. However, pruning wounds can remain susceptible to GTD pathogens for a duration of up to four months, varying based on the specific pathogen (Úrbez-Torres and Gubler 2011; Martinez-Diz et al. 2020; Rosace et al. 2023; Sosnowski et al. 2023).

Ensuring the protection of pruning wounds is crucial for effective management of GTDs, particularly when implemented early in the vineyard’s lifespan (Kaplan et al. 2016; Sosnowski and McCarthy 2017). Traditionally, fungicide treatments have been employed on pruning wounds to prevent infection by GTD pathogens (Moller and Kasimatis 1980; Pearson 1982; Munkvold and Marois 1993; Sosnowski et al. 2008, 2013; Halleen et al. 2010; Rolshausen et al. 2010; Amponsah et al. 2012; Pitt et al. 2012; Sosnowski and Mundy 2019). However, due to environmental concerns regarding the excessive use of synthetic chemicals and their associated risks to human health and the ecosystem, strict regulations have been imposed on synthetic pesticide usage, leading to the removal of the most hazardous chemicals from the market. Consequently, there is a critical need to explore sustainable alternatives for GTD control. One such alternative is biological control, where research has shown that beneficial microorganisms can colonize woody tissues and provide long-lasting broad-spectrum activity against GTD pathogens. The main biological control agent (BCA) used in vineyards to protect pruning wounds is various species of the *Trichoderma* genus. *Trichoderma* primarily acts through mycoparasitism and antibiosis, where it competes with and parasitizes pathogenic fungi while producing antifungal compounds to suppress their growth (Sood et al. 2020). They compete with pathogens for both space and nutrients and can also induce plant resistance (Vinale et al. 2008).

Several studies have demonstrated the effectiveness of *Trichoderma* spp. in protecting pruning wounds against GTD infections (Kotze et al. 2011; Mutawila et al. 2011, 2015, 2016; Reis et al. 2017; Úrbez-Torres et al., 2020; Brown et al. 2021; Blundell and Eskalen 2022; Pollard-Flamand et al. 2022, 2023). Although these studies bring important highlights on the potential of BCAs to protect pruning wounds against GTD infections, the majority of these studies rely on artificial pathogen infections, which does not accurately represent the reality of natural infections. The hypothesis of this study is that the behavior and effectiveness of the evaluated products to protect pruning wounds against trunk disease fungi vary according to the winegrowing region, the growing season, and the group of pathogens present in the vineyard. Therefore, this study aimed to evaluate and compare the efficacy of various liquid formulation fungicide and paste treatments as well as BCAs registered and authorized in Europe against GTDs, for their potential to prevent natural infection of grapevine pruning wounds by trunk disease fungi in two field trials located in France and Spain over three growing seasons.

## Materials and Methods

### Location and characteristics of the experimental vineyards

The assays were carried out at two commercial vineyards located in Samaniego, Álava region (Northern Spain), and Madiran, Nouvelle-Aquitaine region (Southern France), over three years (2020-2023). The vineyard in Samaniego was planted in 2001 (19-years-old) with ‘Tempranillo’ cultivar grafted onto 110 Richter rootstock. On average, Álava region receives around 400-600 mm of rainfall per year and the annual average temperature ranges from approximately 12 to 14 °C. Vines were spaced 1.2 m from center to center, and with an interrow spacing of 2.25 m, trained as bilateral cordons in a trellis system with a spur-pruning (Royat).

The vineyard in Madiran was planted in 1997 (24-years-old) with ‘Cabernet Franc’ cultivar grafted onto SO4 rootstock. The annual average rainfall in this region is 800-1000 mm and the annual average temperature ranges from approximately 12 to 14 °C. Vines were spaced 1.5 m from center to center, and with an interrow spacing of 2.5 m, trained as Guyot system. Standard cultural practices were used in both vineyards during the growing season, and managements of both powdery and downy mildews was performed using only wettable sulphur and copper compounds applied at label dosages and following Integrated Pest Management (IPM) guidelines. Both vineyards were located less than 2 km to an automatic weather station, and its climatic data was considered to be representative.

### Pruning wound protection treatments

Wound protection treatments tested in the present assay are listed in Table 1. We evaluated the efficacy of one chemical and three BCAs formulated products, and also a paste treatment. The chemical and biological products assessed were those commercially formulated and currently registered and available in Europe for control of fungal trunk pathogens. Application rates were selected based on the registered label dosages recommendations. Pyraclostrobin + boscalid (Tessior®) and the paste (Bloccade®) treatments contain a liquid polymer and they already formulated to be directly sprayed to pruning wounds without any previous mixing. Regarding BCA treatments, the conidia viability of both *Trichoderma atroviride* strains (SC1 and I-1237), as well as the mixture *Trichoderma asperellum* ICC012 + *Trichoderma gamsii* ICC080 in the commercial products was tested to be at a minimum of 85% before the assay was set up (Pertot et al. 2016). A serial dilution of the conidia suspension was plated on PDA and the colony-forming units were counted after 24-48 h incubation at room temperature.

**Table 1.**
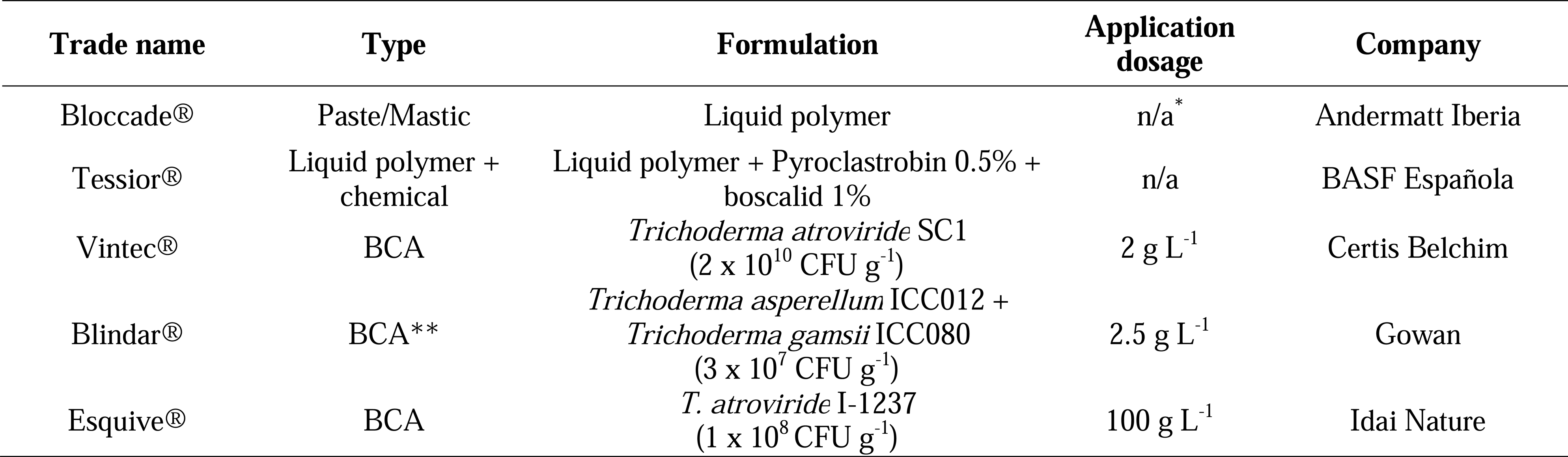
List of products evaluated to protect pruning wounds against fungi associated with grapevine trunk diseases. * n/a, not applicable. ** Biological control agent.

### Field assay and experimental design

On February 21, 2020, in Samaniego, and on March 2, 2020 in Madiran, 1-year-old canes of all vines to be treated were spur-pruned to three buds using secateurs in both vineyards, coinciding with the common pruning time in these regions. Wound treatments were applied by hand until runoff within 2 h after pruning to three wounds per vine. All formulations were applied using a 500 ml hand-held spray bottle with a plastic shield on the nozzle to minimize spray drift. Untreated controls were mock treated with sterile distilled water (SDW). The experiment was set up as a randomized block design with three replicates of ten plants (thirty canes) per wound protectant treatment in each vineyard. Three replicates of ten plants were also used for non-treated controls in each vineyard. The experiment was repeated the following seasons (2021–22 and 2022-23), with pruning and wound treatments applied on 10 February 2021 and 1 February 2022 (Samaniego), and on 3 March 2021 and 17 February 2022 (Madiran).

### Fungal recovery and identification

Canes were harvested from vines above the second bud (about 10 cm long pieces) approximately 12 months after pruning and stored in a 4°C cool room prior to laboratory assessment. Bark was first removed using a sharp knife from each cane. Then, canes were surface sterilised for 1 min in 33% sodium hypochlorite (commercial 40 g Cl/l) and rinsed twice for 1 min each in SDW. After air drying on sterile filter paper to remove moisture excess, each cane was cut into two small pieces (about 12 mm^2^ each) taken from the apical part of the cane to isolate *Trichoderma* spp., and from the margin between discolored or dead and live or apparently healthy wood tissue to isolate GTD pathogens using sterilised secateurs. In both cases, five wood fragments were plated onto each of two plates of Malt Extract Agar (MEA) amended with 0.35 g l^−1^ of streptomycin sulphate (Sigma-Aldrich, St. Louis, MO, USA) (MEAS) giving a total of twenty wood pieces per cane. Cultures were incubated at 25°C under warm fluorescent light for a 12-h photoperiod and inspected daily for 15 days. All growing fungal colonies were transferred to PDA plates for their identification based on molecular methods.

Fungal DNA was extracted from fresh mycelium after 3 weeks of incubation in PDA using the E.Z.N.A. Plant Miniprep Kit (Omega Bio-Tek, Doraville, GA, USA) following manufacturer’s instructions. Initially, the identity of the obtained fungi (pathogens and *Trichoderma* spp.) was revealed through sequencing of the ITS region using the primer pairs ITS1-F (Gardes and Bruns 1993) and ITS4. For specific groups of fungi, additional genes were sequenced. For Botryosphaeriaceae and *Cytospora* spp., the primers EF1-728F and EF1-986R (Carbone and Kohn 1999) were used to amplify part of the translation elongation factor 1-α gene (tef1). The beta-tubulin (tub2) region was amplified using the T1 (O’Donnell and Cigelnik 1997) and Bt2b (Glass and Donaldson 1995) primer set for *Phaeoacremonium* spp., the BTCadF/BTCadR primer set for *Cadophora* spp. (Travadon et al. 2015), or Tub2FD (Aveskamp et al. 2009) and T22 (O’Donnell and Cigelnik 1997) for *Diaporthe* spp. Identitiy of *T. atroviride* SC1 and *T. asperellum* ICC012 + *T. gamsii* ICC080 were revealed by using the specific primers developed by Savazzini et al. (2008) and Gerin et al. (2018), respectively. All PCR products were visualized in 1% agarose gels (agarose D-1 Low EEO, Conda Laboratories) and sequenced in both direction by Eurofins GATC Biotech (Cologne, Germany).

### Data analysis

The average pathogen incidence reduction relative to the control was calculated for the different protectant treatments and years. Statistical analysis was conducted using R, version 4.2.0 (R Core Team 2023). To summarize and visualize the response (pathogen incidence reduction) to the treatments for the different years and experimental vineyard’s locations a clustering analysis and heat map were constructed using the “ComplexHeatmap” package (Gu 2022). The Euclidean distance was used as the similarity measure with complete linkage hierarchical clustering. The incidence data obtained for the different pathogens on each experimental vineyard and for all seasons were analyzed using the Kruskal-Wallis multiple comparison test, followed by the Dunn test with no adjustment for P values using the packages “agricolae” and “dunn.test” (De Mendiburu 2015; R Core Team 2023). The percentage of *Trichoderma* spp. recovery was determined as the mean percentage of infected plants. Significance levels for mean percentages of *Trichoderma* incidence were determined by the Kruskal–Wallis one-way analysis of variance on ranks and mean separation was conducted for Fisher’s least significant differences (LSD) at *P* <0.05.

## Results

### Fungal recovery and identification

The climate data corresponding to the three growing seasons in the two vineyards are detailed in Supp Table 1. In Samaniego, a higher number of isolates associated with GTDs were obtained during the 2022-23 season (2,253 isolates) compared to the 2020-21 season (2,037 isolates) and the 2021-22 season (1,445 isolates) (Table 2). Throughout all seasons, the most prevalent disease was Botryosphaeria dieback, representing 62% of the total isolates in the 2020-21 season, 77.8% in the 2021-22 season, and 61.2% in the 2022-23 season (Table 2). In all seasons, the prevalent fungal species was *Diplodia seriata*, followed by *Botryosphaeria dothidea* in the season 2020-21, *Diplodia mutila* in the season 2021- 22, and *Cytospora viticola* in the season 2022-23 (Supp Table 2).

**Table 2.**
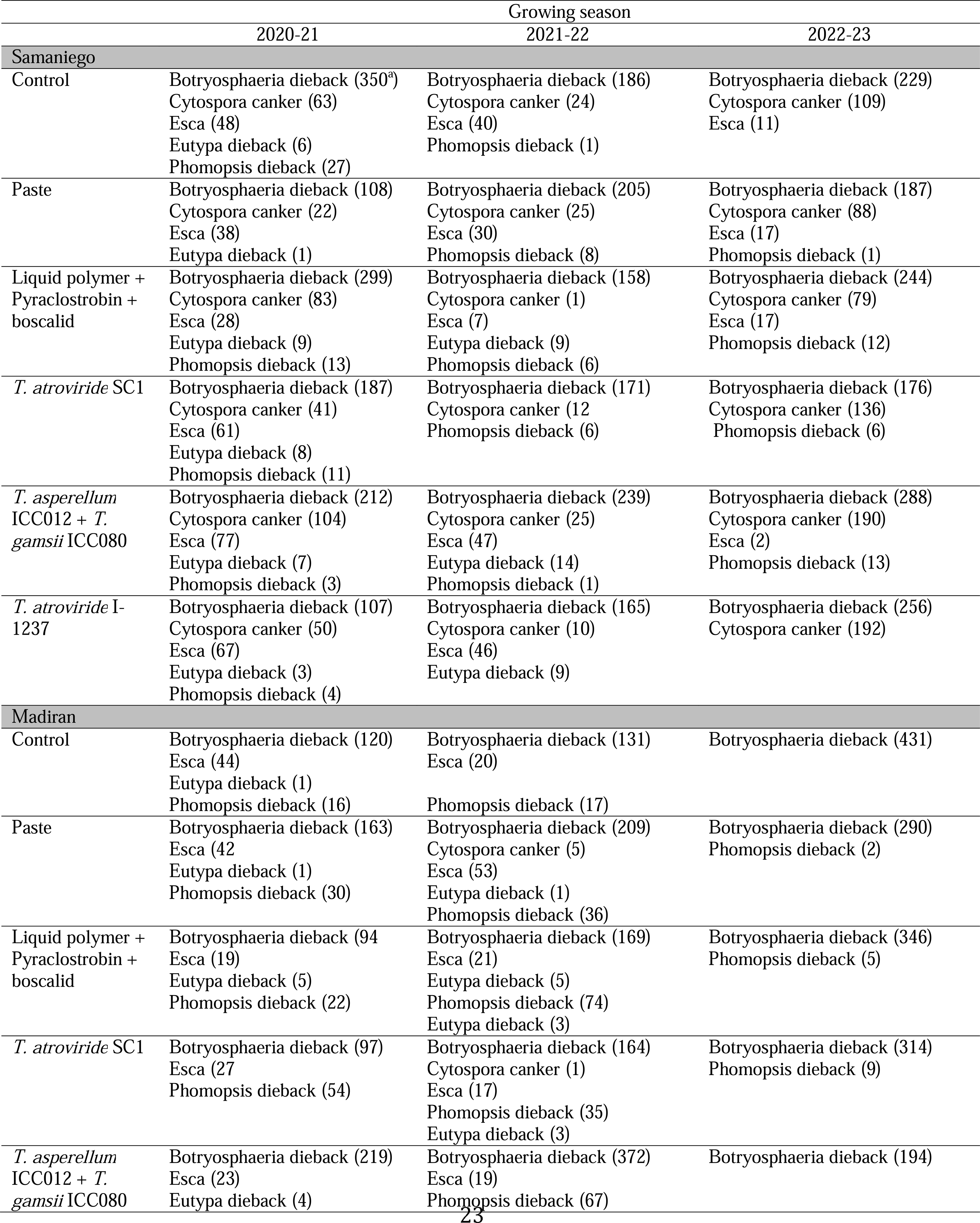
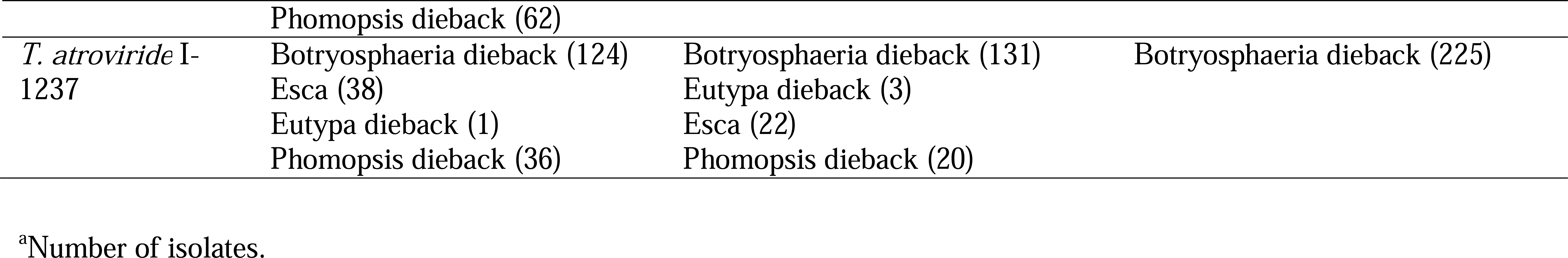
Grapevine trunk diseases detected for each of the treatments during the 3 growing seasons in two vineyards.

In Madiran, a higher number of isolates isolates associated with GTDs were obtained during the 2022-23 season (1,816 isolates) compared to the 2021-22 season (1,598 isolates) and the 2020- 21 season (1,242 isolates) (Table 2). Throughout all seasons, the most prevalent disease was Botryosphaeria dieback, representing 65.8% of the total isolates in the 2020-21 season, 67.3% in the 2021-22 season, and 99.1% in the 2022-23 season (Table 2). In all seasons, the prevalent fungal species was *Diplodia seriata*, followed by *Neofusicoccum* sp. in the seasons 2020-21 and 2022-23, and *Diplodia sapinea* in the season 2021-22 (Supp Table 2).

### Efficacy of evaluated products

Data analysis showed very variable pathogen incidence results among seasons in both experimental vineyards. The heat map and cluster analysis revealed differences in the protectant treatments incidence reduction relative to the control for all seasons in the two experimental vineyards. Three treatments groups (TGs) were distinguished by the cluster analysis (Figure 1). The highest incidence reduction (TG3) for all pathogens was observed for the *T. atroviride* I-1237 treatment in both locations throughout seasons (Figure 1). Pyraclostrobin + boscalid treatments showed intermediate incidence reduction (TG2) for all pathogens in both locations and throughout seasons. Paste and *T. asperellum* ICC012 + *T. gamsii* ICC080 treatments showed high incidence reduction in a single season (2021 and 2023, respectively) but low reduction (TG1) for the other studied periods in the experimental vineyard located in Samaniego (Figure 1).

**Figure 1.**
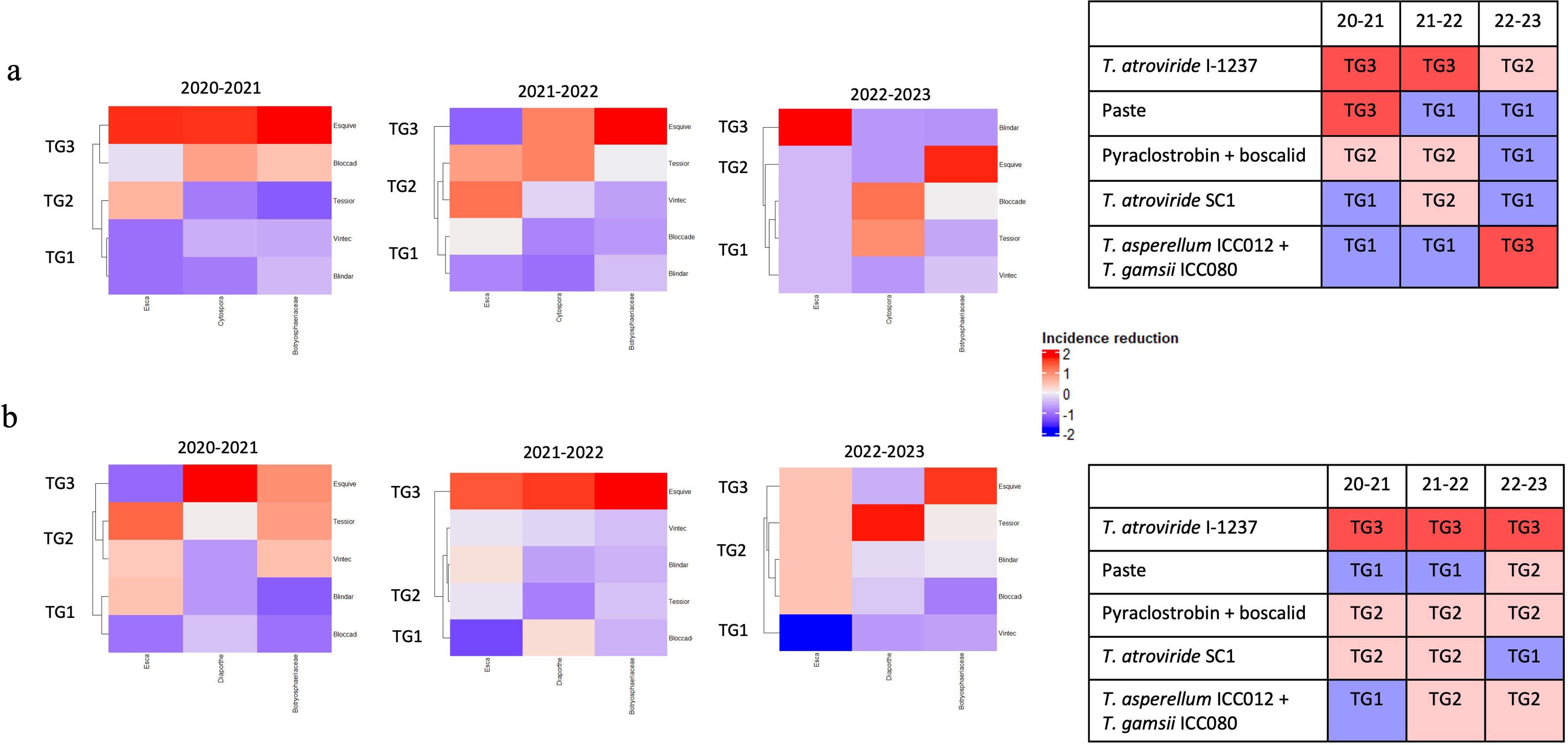

In Samaniego, the multiple comparison analysis showed a significant effect of the treatment (*P*=0.01926) and pathogen (*P*=2.2e-16) on the incidence. *T. atroviride* I-1237 protectant treatment showed a significant reduction (*P*<0.01 and *P*<0.05) in Botryosphaeriaceae spp. incidence relative to the control for all seasons (Figure 2). A significant reduction (*P*<0.05) of Botryosphaeriaceae spp. incidence compared with the control was also observed for the paste treatment but only in season 2020-2021 (Figure 2). In plants treated with *T. atroviride* I-1237 and *T. atroviride* SC1 a significant reduction (*P*<0.05) of esca incidence relative to the control was observed in 2020-2021 and 2021-2022 seasons, respectively (Figure 2). Cytospora canker incidence was reduced (*P*=0.01) compared with the control by *T. atroviride* I-1237 and pyraclostrobin + boscalid treatments in 2021-2022 season (Figure 2).

**Figure 2.**
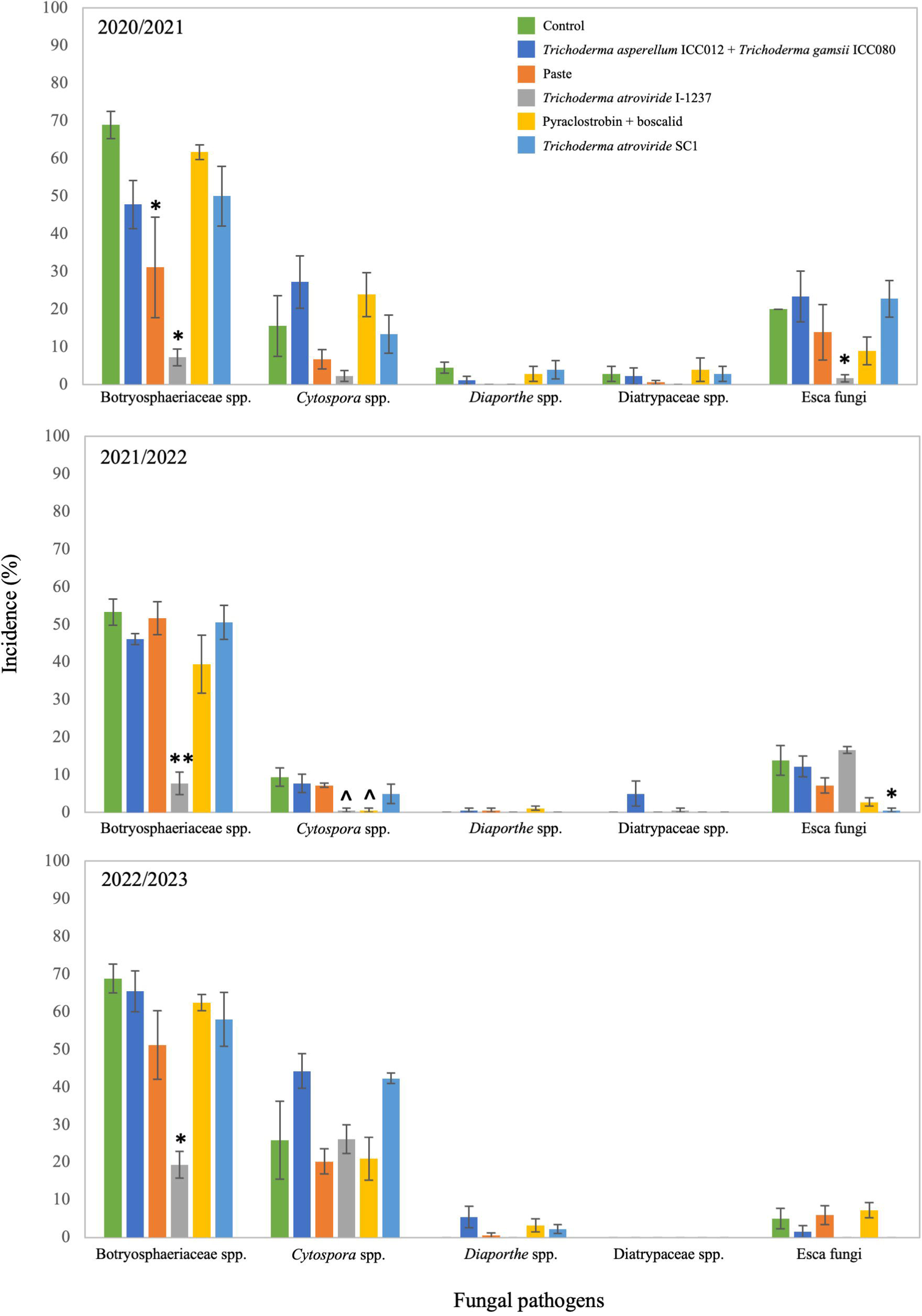

In Madiran, the protectant treatments results showed to be more variable in all seasons for all pathogens. In general, *T. atroviride* I-1237 treatment reduced the Botryosphaeriaceae spp. incidence relative to the control but the effect was significant (*P*<0.01) only in season 2022-2023 (Figure 3). The plants treated with pyraclostrobin + boscalid showed a significant reduction in Botryosphaeriaceae spp. and esca incidence (*P*<0.05 and *P*<0.01, respectively). *Diaporthe* incidence was reduced (*P*=0.01) compared with the control by pyraclostrobin + boscalid treatment in 2022-2023 season (Figure 3).

**Figure 3.**
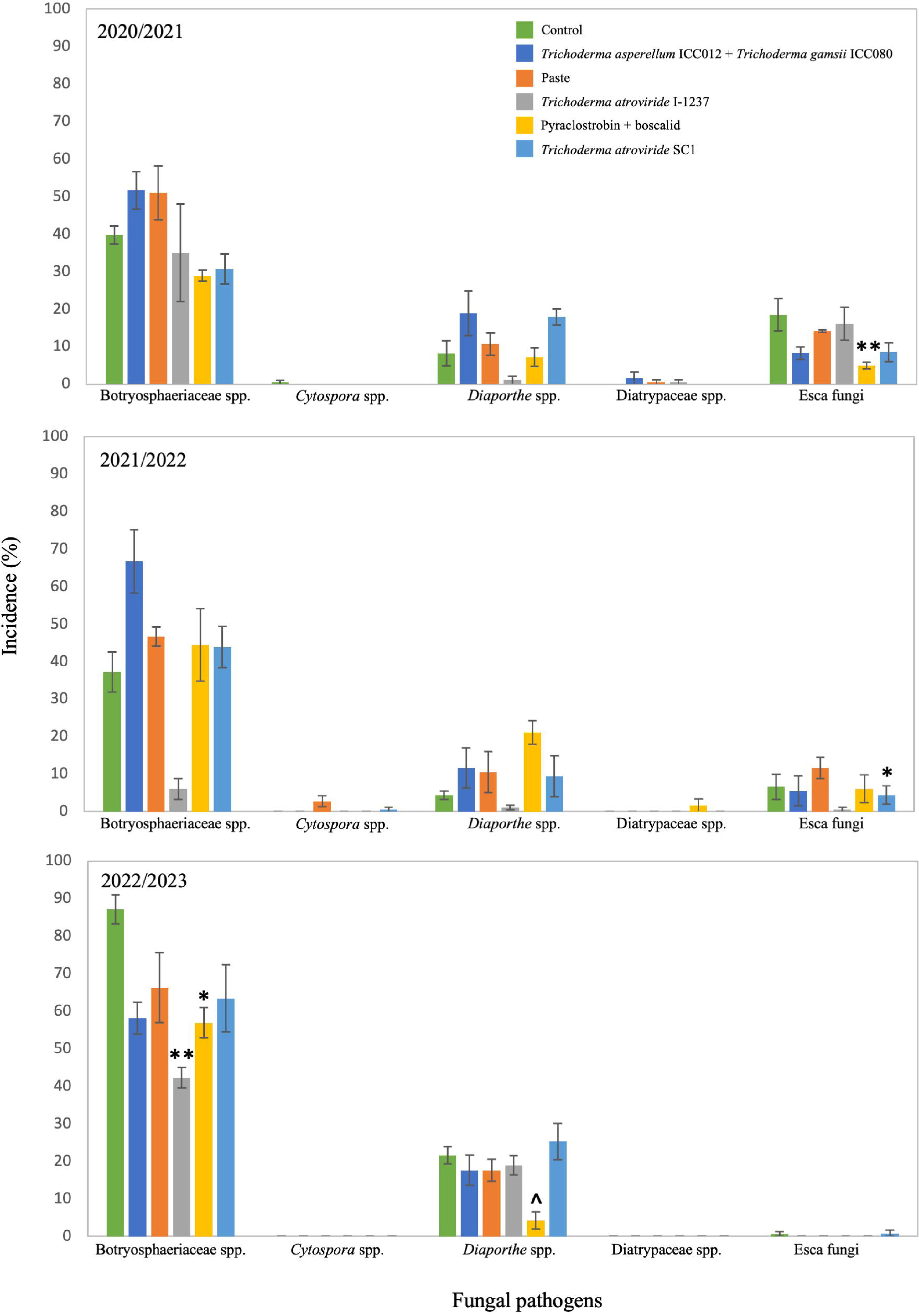

The percentage of isolation of *Trichoderma* spp. is shown in Figure 4. In Samaniego, significant differences in isolation percentage were observed between *T. atroviride* SC1 (67.5%) and *T. asperellum* ICC012 + *T. gamsii* ICC080 (16.7%) in the first seasonof the study. During the second and third growing seasons, the isolation percentages of *T. atroviride* SC1 (93.3% in the second season, 100% in the third season) and *T. atroviride* I-1237 (95% in the second season, 95.2% in the third season) were much higher than that of *T. asperellum* ICC012 + *T. gamsii* ICC080 (48.9% in the second season, 19.4% in the third season), with significant differences in both growing seasons. This same pattern was observed in the first two seasons in Madiran, where significant differences were observed between the isolation percentages of *T. atroviride* SC1 (89.6% in the first season, 94.2% in the second season) and *T. atroviride* I-1237 (85% in the first season, 80.8% in the second season) compared to *T. asperellum* ICC012 + *T. gamsii* ICC080 (32.5% in the first season, 40% in the second season). In the third season of the study, no significant differences were observed between treatments, with low percentages of *Trichoderma* spp. isolation in all cases.

**Figure 4.**
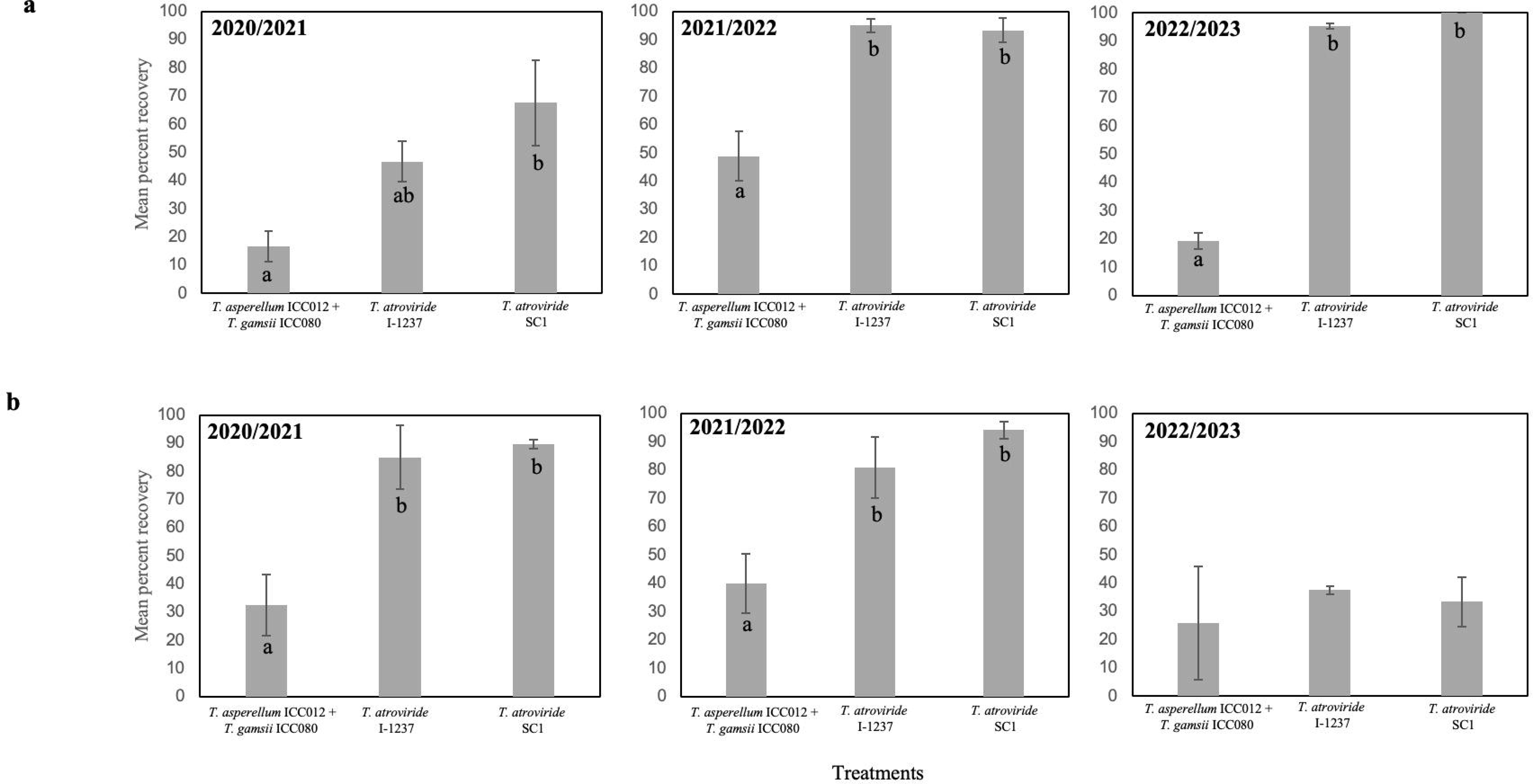

## Discussion

This study is the first comprehensive effort to assess the effectiveness of various preventive treatment approaches, including physical barriers, chemical interventions, and BCAs against natural GTDs infections. This research was conducted in two distinct wine regions over three years. While previous studies have investigated the impact of biological products, such as *Trichoderma*-based treatments, on GTD pathogen infections in the context of natural infection conditions (Halleen et al. 2010; Mutawila et al. 2016; Di Marco et al. 2022), these studies overlooked the potential benefits of physical barriers and chemical treatments.

To date, the major portion of investigations related to protecting pruning wounds has relied on artificial pathogen infections (Amponash et al. 2012; Ayres et al. 2022; Halleen et al. 2010; Martinez-Diz et al. 2021; Mutawila et al. 2016; Reis et al. 2021; Rolshausen et al. 2010; Sosnowski et al. 2008, 2013). This approach was adopted primarily to achieve statistically significant differences, as natural infections often do not reach levels conducive to reliable statistical analysis (Carter and Moller 1971; Ayres et al. 2022). Additionally, there is a prevailing assumption that a product proven effective under conditions of high inoculum pressure from artificially induced trunk disease fungi infections will also exhibit efficacy under the natural infection conditions (Ayres et al. 2022; Carter and Moller 1971). Nevertheless, these previous studies do not take into account the complex interactions among the pathogen and other microorganisms that might be present in the natural environment, which are inherent to field conditions. Furthermore, discarding a product that proves ineffective under conditions of high pathogen inoculum pressure could be misleading, as it might exhibit good performance under conditions of natural infection. For instance, *T. atroviride* I-1237 was found to be ineffective in preventing artificial infections by *D. seriata* (Botryosphaeria dieback) in several vineyards in Galicia (Martínez-Diz et al. 2021), but it has proven effective under conditions of natural infection by Botryosphaeriaceae spp. in the current study.

Although the experiments were conducted in two distinct countries (Spain and France), a common trend emerged in both locations. The prevailing GTD naturally occurring in pruning wounds was Botryosphaeria dieback. Pathogenicity studies have showed that Botryosphaeriaceous species are among the fastest wood-colonizing fungi (Úrbez-Torres et al. 2008), and therefore, the most virulent GTD fungi, which corroborates with the results found in our study. Following Botryosphaeria dieback, the second most prevalent disease in the Samaniego region (Álava, Spain) was Cytospora canker, while in the Madiran region (South of France), Phomopsis dieback, attributed to *Diaporthe* spp., took precedence. Cytospora canker in grapevines remains relatively underexplored, especially in Europe, with the majority of reports on its association with grapevines originating from the United States (Lawrence et al. 2017, 2018; Travadon et al. 2022). The incidence of other common GTDs, such as esca disease and Eutypa dieback (caused by *Diatrypaceae* spp.), was minimal in both experiments. This limited occurrence could be attributed to the slow growth rate of esca-associated pathogens, specifically *Phaeoacremonium* spp. and *Phaeomoniella chlamydospora*, compared to the rapid progression observed in Botryosphaeriaceae pathogens. This distinction can lead to false-negative diagnoses, making the accurate detection of these pathogens challenging, even if they are present in the wood sample.

To mitigate this issue, we isolated pathogens from a substantial number of samples for each treatment, incorporating multiple Petri dish repetitions per sample. This approach enabled us to promptly identify and eliminate fast-growing fungi on a daily basis, ensuring the preservation of fungal viability. Employing molecular techniques, which would entail RNA extraction, was deemed impractical due to its complexity and cost, particularly given the large number of samples involved in our study. This study spanned two vineyards over a three-year period, each subjected to five different treatments alongside a control group, making the use of molecular techniques less feasible in this context.

Environmental factors, including climate conditions, have a substantial impact on the development GTDs and their associated symptoms, as observed in prior studies (Beris et al. 2022; Calvo-Garrido et al. 2021). For instance, esca is thought to be particularly influenced by the prevalence of drought followed by rains in spring and elevated temperatures, which are characteristic of Mediterranean climates (Beris et al. 2022). In contrast, Botryosphaeria dieback infections can manifest in a range of climatic conditions due to the varying climatic requirements of different species responsible for the formation of fruiting structures (Úrbez-Torres 2011). While it might be tempting to attribute the presence of different pathogens in each region to distinct climates, it is worth noting that both regions share the same Köppen climate classification, characterized as a Temperate oceanic climate (cfb). Therefore, it seems less likely that climate alone was responsible for the observed variation in GTD pathogens. An alternative explanation arises from the fact that the majority of GTD pathogens do not exhibit host specificity (Hrycan et al. 2020). This leads us to consider the influence of pathogens present in the surrounding vegetation, which can affect the types of GTD pathogens encountered within a vineyard (Moyo et al. 2019; Hrycan et al. 2020). This explanation appears plausible in understanding why different GTD pathogens are prevalent in two distinct wine regions.

Our findings reveal that none of the products under investigation exhibited complete effectiveness against any of the GTDs. The efficacy of these products was particularly influenced by the specific growing season. A notable exception was observed with the biocontrol agent *T. atroviride* I-1237, which consistently demonstrated effectiveness against Botryosphaeria dieback infections throughout each growing season, irrespective of the location. These outcomes align with prior research highlighting *T. atroviride* I-1237 as a protective agent against Botryosphaeria pathogens (Langa-Lomba et al. 2023; Reis et al. 2017, 2022). It is worth noting, however, that these previous studies primarily employed artificial infections, neglecting the presence of multiple pathogens and variations in concentrations. Therefore, our results complement and confirm that it is an effective treatment against Botryosphaeriaceae spp. infection.

Apart from *T. atroviride* I-1237, the remaining commercial products exhibited efficacy in specific growing seasons or locations against particular diseases. For instance, in Samaniego, the chemical fungicide boscalid + pyraclostrobin demonstrated effectiveness against Cytospora canker and esca in 2022. In Madiran, it was effective against Diaporthe and Botryosphaeria dieback in 2023, as well as esca in 2021 and 2023. Prior research has also emphasized boscalid + pyraclostrobin as a highly effective fungicide against pathogens such as *Diplodia seriata*, *Phaeomoniella chlamydospora*, and *Lasiodiplodia* sp. in grapevine pruning wounds (Martinez-Diz et al. 2021; Reis et al. 2021). However, it is important to note that the efficacy of boscalid + pyraclostrobin, as in our study, was found to be influenced by the specific year in both Martinez-Diz et al. (2021) and Reis et al. (2021). Additional studies examining other pyraclostrobin + boscalid-based fungicides against GTD pathogens in the field have yielded results consistent with these findings, where the fungicide effectiveness was variable depending on the year and location (Brown et al. 2021; Diaz and Latorre 2013).

The treatment showing the least overall effectiveness was the physical barrier (paste), which displayed effectiveness solely against Botryosphaeria pathogens in Madiran during 2021. While pastes are generally considered effective for protecting wounds (Gramaje et al. 2018), they may sometimes yield unsatisfactory results when used without the addition of a fungicidal supplement. As illustrated by Rolshausen and Gubler (2005), the efficacy of a paste in reducing *Eutypa lata* colonization *in vitro*, was only observed when it was supplemented with boric acid. The application of biological commercial products such as *T. asperellum* ICC012 + *T. gamsii* ICC080 and *T*. *atroviride* SC1 yielded inconsistent outcomes, which were contingent on the growing season, location, and the specific disease in question. This variability can be attributed to the utilization of live microorganisms, which can potentially delay disease suppression while amplifying the multitude of factors that influence their success, including soil conditions, pathogen types, and climate (Bale et al. 2008). Both *T. asperellum* ICC012 + *T. gamsii* ICC080 and *T*. *atroviride* SC1 have undergone extensive testing against GTD pathogens in pruning wounds, and they have demonstrated their effectiveness. However, in agreement to our study, their efficacy appears to fluctuate significantly depending on various environmental conditions (Aloi et al. 2015; Bigot et al. 2020; Blundell and Eskalen 2022; Blundell et al. 2020; Di Marco et al. 2022; Martinez-Diz et al. 2021).

In our investigation, the recovery rates of *Trichoderma* spp. in treated plants exhibited significant variability, ranging from 16.7% to 100%. These variations were contingent on several factors, including the specific commercial product used, the growing season, and the geographical location. In general, products based on *T. atroviride* consistently demonstrated higher recovery percentages compared to the *T. asperellum* + *T. gamsii* based-product, owing to differences in the *Trichoderma* strains employed. Prior research has established that the recovery rate can change according to the *Trichoderma* strain used as well as the grapevine cultivar (Mutawila et al. 2011; Carro-Huerga et al. 2021).

Since the BCA application was performed according to manufacturer guidelines, there was non-uniformity in the concentration of *Trichoderma* across the products. Consequently, the *T. asperellum* + *T. gamsii* based-product had a lower *Trichoderma* concentration (3 x 10^7^ cfu/gr) in contrast to *T. atroviride* I-1237 (1 x 10^8^ cfu/gr) and *T. atroviride* SC1 (1 x 10^10^ cfu/gr), which may also account for the observed variations in recovery rates. This discrepancy in recovery percentages was particularly pronounced in the second and third growing seasons in Samaniego. In contrast, in Madiran, during the third growing season, *Trichoderma* recovery rates were noticeably lower than in the preceding two seasons. This discrepancy could be attributed to adverse climate conditions at or subsequent to the BCA application. It is conceivable that abnormal precipitation shortly after the BCA application may have washed away the products, or low temperatures during the year were unfavorable for *Trichoderma* sporulation and colonization. Research by Carro-Huerga et al. (2021) indicated that temperature can impact *Trichoderma*’s colonization capacity, and even though *Trichoderma* can colonize grapevine wood during winter at low temperatures (<10°C), the wood colonization can occasionally be compromised. Martínez-Díaz et al. (2020) discussed how daily mean temperatures of 6.5°C and 8.2°C during the week of *Trichoderma* applications in field grapevines led to poor colonization. Mutawila et al. (2015) recorded temperatures ranging from 12°C to 16°C at the time of inoculation, resulting in a high rate of *Trichoderma* re-isolation from the pruning wounds. Therefore, it is plausible to assume that higher temperatures might positively influence the efficacy of *Trichoderma* colonization.

Building upon prior research and the findings from our study, it is evident that the transition from synthetic chemical fungicides to biological control products offers a multitude of advantages (Mesguida et al. 2023). Chief among these benefits is the absence of residual residues associated with biocontrol products, rendering them exceptionally advantageous later in the growing season. Moreover, these products typically require minimal to no waiting period before harvest. Being classified as ’low-risk substances,’ they possess extended authorization periods and can be safely utilized in environmentally sensitive areas. The increasing utilization of BCAs in plant protection is evidently underway, as indicated by the substantial rise in the use of low-risk EU registered substances in 2022 (Marchand 2023).

Microbial biocontrol agents, owing to their complex mechanisms of action that involve metabolite and enzyme production, resistance induction, resource and space competition, as well as mycoparasitism, pose a significantly reduced risk of inducing pathogen resistance (Leal et al. 2024). Consequently, they assume a vital role in plant protection strategies to mitigate the development of resistance. Furthermore, these biocontrol products represent the exclusive choice for organic production, given their natural origin and minimal environmental impact. They are also ecologically advantageous due to their complete biodegradability, and there is no documented evidence of metabolite bioaccumulation (Pertot et al. 2016). Although some researchers have suggested that microbial biocontrol products may be less effective than synthetic chemicals (Pertot et al. 2016), our study demonstrated their effectiveness as the sole treatment for certain diseases and compared with other chemical-based protectant treatments. For example, *T. atroviride* I-1237 proved effective in reducing infection rates for Botryosphaeria dieback in all seasons and at both locations.

Nevertheless, the broader adoption of microbial biocontrol products faces several constraints in the market. Because these products consist of living organisms, their survival and efficacy are heavily contingent on environmental conditions during and following application, rendering their potential effectiveness variable (Pertot et al. 2016). In this context, it is particularly valuable that some treatments, tested under field conditions in the current study, have proven to be efficient over a period of three growing seaaons and in two different locations. Factors such as exposure to temperatures outside their survival range, UV radiation, and desiccation significantly impact their persistence on plant surfaces. Subsequent to application, the concentration of the microorganism initially remains high but gradually diminishes to levels akin to naturally occurring microorganisms in the specific environment (Savazzini et al. 2009). In many instances, this duration proves insufficient to reduce pathogen inoculum to a safe level. Additionally, the necessary concentration threshold for achieving effectiveness can render these treatments cost-prohibitive for growers, making them economically unsustainable (Pertot et al. 2016). It is important to emphasize that in our study, the persistence of *T. atroviride*-based formulations was significant, as evidenced by the high recovery percentages obtained.

To conclude, this study underscores the importance of customizing treatments for specific diseases, taking into account the influence of environmental factors. This is especially pertinent when considering BCAs. Further research is warranted to expand our understanding of these microorganisms in diverse grape growing regions with differing climate conditions and grapevine genotypes, considering the substantial variability in results.

## Acknowledgements

This research was financially supported by the Project EFA 324/19 – VITES QUALITAS, which has been 65% co-financed by the European Regional Development Fund (ERDF) through the Interreg V-A Spain-France-Andorra program (POCTEFA 2014-2020).

## Notes

### Competing Interest Statement

The authors have declared no competing interest.

## Literature Cited

1. Aloi, C., Reggiori, F., Bigot, G., Montermini, P., Bortolotti, R., Nannini, F. et al. 2015. REMEDIER® (*Trichoderma asperellum* and *Trichoderma gamsii*): a new opportunity to control the esca disease complex. Five years of results of field trials in Italy. Phytopathol. Mediterr. 54:430–431.

2. Amponsah, N. T., Jones, E., Ridgway, H. J., and Jaspers, M. V. 2012. Evaluation of fungicides for the management of Botryosphaeria dieback diseases of grapevines. Pest Manag. Sci. 68(5):676–683.

3. Aveskamp, M. M., Verkley, G. J. M., De Gruyter, J., Murace, M., Perelló, A., Woundenberg, J., et al. 2009. DNA phylogeny reveals polyphyly of Phoma section *Peyronellaea* and multiple taxonomic novelties. Mycologia 101:363–382.

4. Ayres, M. R., Billones-Baaijens, R., Savocchia, S., Scott, E. S., and Sosnowski, M. R. 2022. Critical timing of fungicide application for pruning wound protection to control grapevine trunk diseases. Aust. J. Grape Wine Res. 28(1):70–74.

5. Bale, J. S., Van Lenteren, J. C., and Bigler, F. 2008. Biological control and sustainable food production. Philos. Trans. R. Soc. B: Biol. 363(1492):761–776.

6. Baloyi, M. A., Halleen, F., Mostert, L., and Eskalen, A. 2016. First report of *Phaeomoniella chlamydospora* pycnidia as Petri disease inoculum sources in South African vineyards. Plant Dis. 100(12):2528–2528.

7. Beris, E., Selim, M., Kechagia, D., and Evangelou, A. 2022. Overview of the Esca Complex as an Increasing Threat in Vineyards Worldwide: Climate Change, Control Approaches and Impact on Grape and Wine Quality. In Recent Advances in Grapes and Wine Production-New Perspectives for Quality Improvement. IntechOpen.

8. Bertsch, C., Ramírez-Suero, M., Magnin-Robert, M., Larignon, P., Chong, J., Abou-Mansour, E., et al. 2013. Grapevine trunk diseases: complex and still poorly understood. Plant Pathol. 62(2):243–265.

9. Bigot, G., Sivilotti, P., Stecchina, M., Lujan, C., Freccero, A., and Mosetti, D. 2020. Long-term effects of *Trichoderma asperellum* and *Trichoderma gamsii* on the prevention of esca in different vineyards of Northeastern Italy. Crop Protect. 137:105–264.

10. Blundell, R., Lynch, M., Haden, T., Arreguin, M., Gallagher, T., and Eskalen, A. 2020. Evaluation of Vintec (*Trichoderma atroviride* SC1) as Pruning Wound Protectants Against Selected Fungi Associated with Grapevine Trunk Diseases.

11. Blundell, R., and Eskalen, A. 2022. Biological and chemical pruning wound protectants reduce infection of grapevine trunk disease pathogens. Cal. Agric. 75(3):128–134.

12. Brown, A. A., Travadon, R., Lawrence, D. P., Torres, G., Zhuang, G., and Baumgartner, K. 2021. Pruning-wound protectants for trunk-disease management in California table grapes. Crop Protect. 141:105490.

13. Bruez, E., Lecomte, P., Grosman, J., Doublet, B., Bertsch, C., Fontaine, F., et al. 2013. Overview of grapevine trunk diseases in France in the 2000s. Phytopathol. Mediterr. 262–275.

14. Calvo-Garrido, C., Songy, A., Marmol, A., Roda, R., Clément, C., and Fontaine, F. 2021. Description of the relationship between trunk disease expression and meteorological conditions, irrigation and physiological response in Chardonnay grapevines. OENO One 55(2):97–113.

15. Carbone, I. and Kohn, L.M. 1999. A method for designing primer sets for speciation studies in filamentous ascomycetes. Mycologia 91:553–556.

16. Carro-Huerga, G., Mayo-Prieto, S., Rodríguez-González, Á., Álvarez-García, S., Gutiérrez, S., and Casquero, P. A. 2021. The influence of temperature on the growth, sporulation, colonization, and survival of *Trichoderma* spp. in grapevine pruning wounds. Agronomy 11(9):1771.

17. Carter, M. V., and Moller, W. J. 1971. The quantity of inoculum required to infect apricot and other Prunus species with *Eutypa armeniacae*. Aust. J. Exp. Agric. 11(53):684–686.

18. De Mendiburu, F. 2020. Agricolae: Statistical Procedures for Agricultural Research. https://CRAN.R-project.org/package=agricolae

19. Díaz, G. A., and Latorre, B. A. 2013. Efficacy of paste and liquid fungicide formulations to protect pruning wounds against pathogens associated with grapevine trunk diseases in Chile. Crop Protec. 46:106–112.

20. Di Marco, S., Metruccio, E. G., Moretti, S., Nocentini, M., Carella, G., Pacetti, A., et al. 2022. Activity of *Trichoderma asperellum* strain ICC 012 and *Trichoderma gamsii* strain ICC 080 toward diseases of esca complex and associated pathogens. Front. Microbiol. 12:813410.

21. Eskalen, A., and Gubler, W. D. 2001. Association of spores of *Phaeomoniella chlamydospora*, Phaeoacremonium inflatipes, and Pm. aleophilum with grapevine cordons in California. Phytopathol. Mediterr. S429–S432.

22. Gardes M and Bruns TD. 1993. ITS primers with enhanced specificity for basidiomycetes: application to the identification ofmycorrhizae and rusts. Mol. Ecol. 2:113–118.

23. Gendloff, E. H., Ramsdell, D. C., and Burton, C. L. 1983. Fungicidal control of *Eutypa armeniacae* infecting Concord grapevine in Michigan. Plant Dis. 67(7):754–756.

24. Gerin, D., Pollastro, S., Raguseo, C., Angelini, R. M. and Faretra, F. 2018. A Ready-to-Use Singleand Duplex-TaqMan-qPCR Assay to Detect and Quantify the Biocontrol Agents *Trichoderma asperellum* and *Trichoderma gamsii*. Front. Microbiol. 9:2073.

25. Glass, N. L. and Donaldson, G. C. 1995. Development of primer sets designed for use with the PCR to amplify conserved genes from filamentous infection due to *Phaeoacremonium* spp. J. Clin. Microbiol. 41:1332–1336.

26. González-Domínguez, E., Caffi, T., Languasco, L., Latinovic, N., Latinovic, J., and Rossi, V. 2021. Dynamics of *Diaporthe ampelina* conidia released from grape canes that overwintered in the vineyard. Plant Dis. 105(10):3092–3100.

27. Gramaje, D., Urbez-Torres, J. R., and Sosnowski, M. R. 2018. Managing grapevine trunk diseases with respect to etiology and epidemiology: current strategies and future prospects. Plant Dis. 102(1):12–39.

28. Gu, Z. 2022. Complex Heatmap Visualization, iMeta. DOI:10.1002/imt2.43.

29. Halleen, F., Fourie, P. H., and Lombard, P. J. 2010. Protection of grapevine pruning wounds against *Eutypa lata* by biological and chemical methods. S. Afr. J. Enol. Vitic. 31(2):125–132.

30. Hrycan, J., Hart, M., Bowen, P., Forge, T., and Urbez-Torres, J. R. 2020. Grapevine trunk disease fungi: Their roles as latent pathogens and stress factors that favour disease development and symptom expression. Phytopathol. Mediterr. 59(3):395–424.

31. Kaplan, J., Travadon, R., Cooper, M., Hillis, V., Lubell, M., and Baumgartner, K. 2016. Identifying economic hurdles to early adoption of preventative practices: The case of trunk diseases in California winegrape vineyards. Wine Econ. Pol. 5(2):127–141.

32. Kotze, C., Van Niekerk, J., Mostert, L., Halleen, F., and Fourie, P. 2011. Evaluation of biocontrol agents for grapevine pruning wound protection against trunk pathogen infection. Phytopathol. Mediterr. 50:S247–S263.

33. Kuntzmann, P., Villaume, S., amd Bertsch, C. 2009. Conidia dispersal of Diplodia species in a French vineyard. Phytopathol. Mediterr. 48(1):150–154.

34. Langa-Lomba, N., González-García, V., Venturini-Crespo, M. E., Casanova-Gascón, J., Barriuso-Vargas, J. J., and Martín-Ramos, P. 2023. Comparison of the Efficacy of Trichoderma and Bacillus Strains and Commercial Biocontrol Products against Grapevine Botryosphaeria Dieback Pathogens. Agronomy 13(2):533.

35. Larignon, P., and Dubos, B. 2000. Preliminary studies on the biology of Phaeoacremonium (*Vitis vinifera* L.-esca disease). Phytopathol. Mediterr. (Italy):39(1).

36. Lawrence, D. P., Travadon, R., Pouzoulet, J., Rolshausen, P. E., Wilcox, W. F., and Baumgartner, K. 2017. Characterization of Cytospora isolates from wood cankers of declining grapevine in North America, with the descriptions of two new Cytospora species. Plant Pathol. 66(5):713–725.

37. Lawrence, D. P., Travadon, R., and Baumgartner, K. 2018. Novel Seimatosporium species from grapevine in northern California and their interactions with fungal pathogens involved in the trunk-disease complex. Plant Dis. 102(6):1081–1092.

38. Leal, C., Eichmeier, A., Stuskova, K., Armengol, J., Bujanda, R., Fontaine, F., et al. 2024. Biocontrol agents establishment and their impact on rhizosphere microbiome and induced grapevine defenses is highly soil-dependent. Phytobiomes.

39. Marchand, P.A. 2023. Evolution of plant protection active substances in Europe: the disappearance of chemicals in favour of biocontrol agents. Environ. Sci. Pollut. Res. 30:1–17.

40. Martínez-Diz, P., Eichmeier, A., Spetik, M., Bujanda, R., Díaz-Fernández, Á., Díaz-Losada, E., and Gramaje, D. 2020. Grapevine pruning time affects natural wound colonization by wood- invading fungi. Fung. Ecol. 48:100994.

41. Martínez-Diz, M., Díaz-Losada, E., Díaz-Fernández, Á., Bouzas-Cid, Y., and Gramaje, D. 2021. Protection of grapevine pruning wounds against *Phaeomoniella chlamydospora* and *Diplodia seriata* by commercial biological and chemical methods. Crop Protect. 143:105465.

42. Mesguida, O., Haidar, R., Yacoub, A., Dreux-Zigha, A., Berthon, J.-Y., Guyoneaud, R., et al. 2023. Microbial Biological Control of Fungi Associated with Grapevine Trunk Diseases: A Review of Strain Diversity, Modes of Action, and Advantages and Limits of Current Strategies. J. Fungi 9:638.

43. Moller, W. J., and Kasimatis, A. N. 1980. Protection of grapevine pruning wounds from Eutypa dieback. Plant Dis. 64(3):278–280.

44. Moyo, P., Mostert, L., and Halleen, F. 2019. Diatrypaceae species overlap between vineyards and natural ecosystems in South Africa. Fungal Ecol. 39:142–151.

45. Munkvold, G. P., and Marois, J. J. 1993. Efficacy of natural epiphytes and colonizers of grapevine pruning wounds for biological control of Eutypa dieback. Phytopathol. 83(6):624–629.

46. Murolo, S., and Romanazzi, G. 2014. Effects of grapevine cultivar, rootstock and clone on esca disease. Australas. Plant Pathol. 43, 215–221.

47. Mutawila, C., Fourie, P. H., Halleen, F., and Mostert, L. 2011. Grapevine cultivar variation to pruning wound protection by Trichoderma species against trunk pathogens. Phytopathol. Mediterr. 50:S264–S276.

48. Mutawila, C., Halleen, F., and Mostert, L. 2016. Optimisation of time of application of Trichoderma biocontrol agents for protection of grapevine pruning wounds. Aust. J. Grape Wine Res. 22(2):279–287.

49. Mutawila, C., Halleen, F., and Mostert, L. 2015. Development of benzimidazole resistant Trichoderma strains for the integration of chemical and biocontrol methods of grapevine pruning wound protection. BioControl 60:387–399.

50. O’Donnell, K., Cigelnik, E., 1997. Two divergent intragenomic rDNA ITS2 types within a monophyletic lineage of the fungus Fusarium are nonorthologous. Mol. Phylogenet. Evol. 7:103–116.

51. Pearson, R. C. 1982. Protection of grapevine pruning wounds from infection by *Eutypa armeniacae* in New York State. Am. J. Enol. Vit. 33(1):51–52.

52. Pertot, I., Prodorutti, D., Colombini, A., and Pasini, L. 2016. *Trichoderma atroviride* SC1 prevents *Phaeomoniella chlamydospora* and *Phaeoacremonium aleophilum* infection of grapevine plants during the grafting process in nurseries. BioControl 61:257–267.

53. Pitt, W. M., Sosnowski, M. R., Huang, R., Qiu, Y., Steel, C. C., and Savocchia, S. 2012. Evaluation of fungicides for the management of Botryosphaeria canker of grapevines. Plant Dis. 96(9):1303–1308.

54. Pollard-Flamand, J., Boulé, J., Hart, M., and Úrbez-Torres, J.R. 2022. Biocontrol activity of *Trichoderma* species isolated from grapevines in British Columbia against Botryosphaeria dieback fungal pathogens. J. Fungi 8:409.

55. Pollard-Flamand, J., Boulé, J., Hart, M., and Úrbez-Torres, J.R. 2023. Biological control of Botryosphaeria dieback of grapevine in British Columbia, Canada. Am. J. Enol. Vitic. 74:0740034.

56. R Core Team 2023. R: A Language and Environment for Statistical Computing. R Foundation for Statistical Computing, Vienna, Austria. https://www.R-project.org/

57. Reis, P., Gaspar, A., Alves, A., Fontaine, F., and Rego, C. 2021. Combining an HA+ Cu (II) site-targeted copper-based product with a pruning wound protection program to prevent infection with *Lasiodiplodia* spp. in grapevine. Plants 10(11):2376.

58. Reis, P., Gaspar, A., Alves, A., Fontaine, F., and Rego, C. 2022. Response of different grapevine cultivars to infection by *Lasiodiplodia theobromae* and *Lasiodiplodia mediterranea*. Plant Dis. 106(5):1350–1357.

59. Reis, P., Letousey, P., and Rego, C. 2017. *Trichoderma atroviride* strain I-1237 protects pruning wounds against grapevine wood pathogens. Phytopathol. Mediterr. 56:580.

60. Rolshausen, P. E., and Gubler, W. D. 2005. Use of boron for the control of Eutypa dieback of grapevines. Plant Dis. 89(7):734–738.

61. Rolshausen, P. E., Úrbez-Torres, J. R., Rooney-Latham, S., Eskalen, A., Smith, R. J., and Gubler, W. D. 2010. Evaluation of pruning wound susceptibility and protection against fungi associated with grapevine trunk diseases. Am. J. Enol. Vit. 61(1):113–119.

62. Rosace, M. C., Legler, S. E., Salotti, I., and Rossi, V. 2023. Susceptibility of pruning wounds to grapevine trunk diseases: A quantitative analysis of literature data. Front. Plant Sci. 14:1063932.

63. Savazzini, F., Longa, C. M. O., and Pertot, I. 2009. Impact of the biocontrol agent *Trichoderma atroviride* SC1 on soil microbial communities of a vineyard in northern Italy. Soil Biol. Biochem. 41(7):1457–1465.

64. Savazzini, F., Longa, C. M. O., Pertot, I., and Gessler, C. 2008. Real-time PCR for detection and quantification of the biocontrol agent *Trichoderma atroviride* strain SC1 in soil. J. Microbiol. Methods 73(2):185–194.

65. Sood, M., Kapoor, D., Kumar, V., Sheteiwy, M. S., Ramakrishnan, M., Landi, M., et al. 2020. Trichoderma: The “secrets” of a multitalented biocontrol agent. Plants 9(6):762.

66. Sosnowski, M. R., Ayres, M. R., Billones-Baaijens, R., Savocchia, S., and Scott, E. S. 2023. Susceptibility of pruning wounds to grapevine trunk disease pathogens *Eutypa lata* and *Diplodia seriata* in three climatic conditions in Australia. Fungal Ecol. 64:101260.

67. Sosnowski, M. R., Luque, J., Loschiavo, A. P., Martos, S., Garcia-Figueres, F., Wicks, T. J., and Scott, E. S. 2011. Studies on the effect of water and temperature stress on grapevines inoculated with Eutypa lata. Phytopathol. Mediterr. 50:S127–S138.

68. Sosnowski, M. R., Creaser, M. L., Wicks, T. J., Lardner, R., and Scott, E. S. 2008. Protection of grapevine pruning wounds from infection by *Eutypa lata*. Aust. J. Grape Wine Res. 14(2):134–142.

69. Sosnowski, M. R., Loschiavo, A. P., Wicks, T. J., and Scott, E. S. 2013. Evaluating treatments and spray application for the protection of grapevine pruning wounds from infection by *Eutypa lata*. Plant Dis. 97(12):1599–1604.

70. Sosnowski, M. R., and Mundy, D. C. 2019. Pruning wound protection strategies for simultaneous control of Eutypa and Botryosphaeria dieback in New Zealand. Plant Dis. 103(3):519–525.

71. Sosnowski, M., and McCarthy, G. 2017. Trunk disease: economic impact of grapevine trunk disease management in Sauvignon Blanc vineyards of New Zealand. Wine Vit. J. 32(5).

72. Travadon, R., Lawrence, D.P., Rooney-Latham, S., Gubler, W.D., Wilcox, W.W., Rolshausen, P.E., and Baumgartner, K. 2015. Cadophora species associated with wood decay of grapevine in North America. Fungal Biol. 119:53–66.

73. Travadon, R., Lawrence, D. P., Moyer, M. M., Fujiyoshi, P. T., and Baumgartner, K. 2022. Fungal species associated with grapevine trunk diseases in Washington wine grapes and California table grapes, with novelties in the genera *Cadophora*, Cytospora, and Sporocadus. Front. Fung. Biol. 3:1018140.

74. Úrbez-Torres, J. R. 2011. The status of Botryosphaeriaceae species infecting grapevines. Phytopathol. Mediterr. 50:S5–S45.

75. Úrbez-Torres, J. R., Leavitt, G. M., Guerrero, J. C., Guevara, J., and Gubler, W. D. 2008. Identification and pathogenicity of *Lasiodiplodia theobromae* and *Diplodia seriata*, the causal agents of bot canker disease of grapevines in Mexico. Plant Dis. 92(4):519–529.

76. Úrbez-Torres, J. R., and Gubler, W. D. 2011. Susceptibility of grapevine pruning wounds to infection by *Lasiodiplodia theobromae* and *Neofusicoccum parvum*. Plant Pathol. 60(2):261–270.

77. Úrbez-Torres, J. R., Bruez, E., Hurtado, J., and Gubler, W. D. 2010. Effect of temperature on conidial germination of Botryosphaeriaceae species infecting grapevines. Plant Dis. 94(12):1476–1484.

78. Úrbez-Torres, J. R., Tomaselli, E., Pollard-Flamand, J., Boule, J., Gerin, D., and Pollastro, S. (2020). Characterization of *Trichoderma* isolates from southern Italy, and their potential biocontrol activity against grapevine trunk disease fungi. Phytopatho. Mediterr. 59(3):425–439.

79. Valencia, D., Torres, C., Camps, R., Lopez, E., Celis-Diez, J. L., and Besoain, X. 2015. Dissemination of Botryosphaeriaceae conidia in vineyards in the semiarid Mediterranean climate of the Valparaíso Region of Chile. Phytopatho. Mediterr. 394–402.

80. Van Niekerk, J. M., Calitz, F. J., Halleen, F., and Fourie, P. H. 2010. Temporal spore dispersal patterns of grapevine trunk pathogens in South Africa. Eur. J. Plant Pathol. 127:375–390.

81. Vinale, F., Sivasithamparam, K., Ghisalberti, E. L., Marra, R., Woo, S. L., and Lorito, M. 2008. Trichoderma–plant–pathogen interactions. Soil Biol. Biochem. 40(1):1–10.

