## Supplementary tables for "Evaluating treatments for the protection of grapevine pruning wounds from natural infection by trunk disease fungi"

|  | 2020-21 season | | 2021-22 season | | 2022-23 season | |
| --- | --- | --- | --- | --- | --- | --- |
| Samaniego |  |  |  |  |  |  |
|  | Week of the experiment | Month of the experiment | Week of the experiment | Month of the experiment | Week of the experiment | Month of the experiment |
| Daily mean temperature (ºC) | 9.1 | 9.1 | 7.9 | 8.6 | 5.5 | 7.1 |
| Relative humidity (%) | 67.5 | 76.5 | 85.7 | 82.5 | 79.7 | 74.9 |
| Rain events of > 1 mm | 0 | 0 | 1 | 7 | 1 | 1 |
| Precipitation (mm) | - | 4 | 8.4 | 57.8 | 1.6 | 1.6 |
| Madiran |  |  |  |  |  |  |
| Daily mean temperatura (ºC) | 9.6 | 10.3 | 9.2 | 10.5 | 10.9 | 9.2 |
| Relative humidity (%) | 88.9 | 83.2 | 78.5 | 73.3 | 84.1 | 88.3 |
| Rain events of > 1 mm | 5 | 12 | 2 | 8 | 3 | 10 |
| Precipitation (mm) | 49.5 | 121 | 4 | 14 | 11.6 | 46.3 |

**Supplementary Table 1.** Climate data of the three growing seasons in the two vineyards.

**Supplementary Table 2.** Grapevine trunk diseases and fungal species detected for each of the treatments during the three growing seasons in two vineyards.

|  | **Growing season** | | |
| --- | --- | --- | --- |
|  | **2020-21** | **2021-22** | **2022-23** |
| **Samaniego** |  |  |  |
| **Control** | **Botryosphaeria dieback (350^a^)**  *Botryosphaeria dothidea* (98)  *Diplodia mutila* (72)  *Diplodia seriata* (180)  **Cytospora canker (63)**  *Cytospora* sp. (40)  *Cytospora viticola* (23)  **Esca (48)**  *Phaeoacremonium* sp. (1)  *Phaeomoniella chlamydospora* (47)  **Eutypa dieback (6)**  *Eutypa lata* (6)  **Phomopsis dieback (27)**  *Diaporthe* sp. (26)  *Diaporthe rudis* (1) | **Botryosphaeria dieback (186)**  *Diplodia mutila* (8)  *Diplodia seriata* (178)  **Cytospora canker (24)**  *Cytospora* sp. (5)  *Cytospora viticola* (19)  **Esca (40)**  *Phaeomoniella chlamydospora* (40)  **Phomopsis dieback (1)**  *Diaporthe* *ampelina* (1) | **Botryosphaeria dieback (229)**  *Diplodia mutila* (17)  *Diplodia seriata* (212)  **Cytospora canker (109)**  *Cytospora* *rhodophila* (1)  *Cytospora viticola* (108)  **Esca (11)**  *Phaeomoniella chlamydospora* (11) |
| **Vintec** | **Botryosphaeria dieback (187)**  *Botryosphaeria dothidea* (76)  *Diplodia mutila* (83)  *Diplodia seriata* (28)  **Cytospora canker (41)**  *Cytospora* sp. (6)  *Cytospora viticola* (35)  **Esca (61)**  *Phaeoacremonium* sp. (7)  *Phaeomoniella chlamydospora* (54)  **Eutypa dieback (8)**  *Eutypa lata* (8)  **Phomopsis dieback (11)**  *Diaporthe ampelina* (9)  *Diaporthe* sp. (2) | **Botryosphaeria dieback (171)**  *Botryosphaeria dothidea* (12)  *Diplodia mutila* (21)  *Diplodia seriata* (138)  **Cytospora canker (12)**  *Cytospora* sp. (1)  *Cytospora viticola* (11)  **Phomopsis dieback (6)**  *Diaporthe ampelina* (5)  *Diaporthe* *viticola* (1) | **Botryosphaeria dieback (176)**  *Diplodia mutila* (24)  *Diplodia seriata* (152)  **Cytospora canker (136)**  *Cytospora viticola* (136)  **Phomopsis dieback (6)**  *Diaporthe ampelina* (5)  *Diaporthe* sp. (1) |
| **Tessior** | **Botryosphaeria dieback (299)**  *Botryosphaeria dothidea* (99)  *Diplodia mutila* (71)  *Diplodia seriata* (128)  *Neofusicoccum parvum* (1)  **Cytospora canker (83)**  *Cytospora predappioensis* (1)  *Cytospora* sp. (50)  *Cytospora viticola* (32)  **Esca (28)**  *Phaeoacremonium* sp. (1)  *Phaeomoniella chlamydospora* (27)  **Eutypa dieback (9)**  *Eutypa lata* (9)  **Phomopsis dieback (13)**  *Diaporthe* sp. (13) | **Botryosphaeria dieback (158)**  *Diplodia mutila* (20)  *Diplodia seriata* (138)  **Cytospora canker (1)**  *Cytospora viticola* (1)  **Esca (7)**  *Phaeomoniella chlamydospora* (7)  **Eutypa dieback (9)**  *Eutypa lata* (9)  **Phomopsis dieback (6)**  *Diaporthe ampelina* (1)  *Diaporthe* *rudis* (2)  *Diaporthe* sp. (1)  *Diaporthe* *viticola* (2) | **Botryosphaeria dieback (244)**  *Botryosphaeria dothidea* (16)  *Diplodia seriata* (228)  **Cytospora canker (79)**  *Cytospora predappioensis* (1)  *Cytospora* *rhodophila* (9)  *Cytospora viticola* (69)  **Esca (17)**  *Cadophora* sp. (1)  *Phaeomoniella chlamydospora* (16)  **Phomopsis dieback (12)**  *Diaporthe* *rudis* (9)  *Diaporthe* sp. (3) |
| **Bloccade** | **Botryosphaeria dieback (108)**  *Botryosphaeria dothidea* (84)  *Diplodia mutila* (6)  *Diplodia seriata* (18)  **Cytospora canker (22)**  *Cytospora predappioensis* (1)  *Cytospora viticola* (21)  **Esca (38)**  *Phaeoacremonium* *minimum* (1)  *Phaeoacremonium* sp. (5)  *Phaeomoniella chlamydospora* (32)  **Eutypa dieback (1)**  *Eutypa lata* (1) | **Botryosphaeria dieback (205)**  *Botryosphaeria dothidea* (21)  *Diplodia mutila* (13)  *Diplodia seriata* (171)  **Cytospora canker (25)**  *Cytospora sacculus* (1)  *Cytospora* sp. (3)  *Cytospora viticola* (21)  **Esca (30)**  *Phaeoacremonium* *minimum* (2)  *Phaeomoniella chlamydospora* (28)  **Phomopsis dieback (8)**  *Diaporthe* *rudis* (2)  *Diaporthe* *viticola* (6) | **Botryosphaeria dieback (187)**  *Diplodia seriata* (165)  *Diplodia mutila* (22)  **Cytospora canker (88)**  *Cytospora viticola* (88)  **Esca (17)**  *Cadophora* sp. (1)  *Phaeomoniella chlamydospora* (16)  **Phomopsis dieback (1)**  *Diaporthe* *ampelina* (1) |
| **Blindar** | **Botryosphaeria dieback (212)**  *Botryosphaeria dothidea* (1)  *Diplodia mutila* (49)  *Diplodia seriata* (162)  **Cytospora canker (104)**  *Cytospora predappioensis* (1)  *Cytospora viticola* (93)  *Cytospora* sp. (10)  **Esca (77)**  *Cadophora* sp. (3)  *Phaeomoniella chlamydospora* (74)  **Eutypa dieback (7)**  *Eutypa lata* (7)  **Phomopsis dieback (3)**  *Diaporthe ampelina* (3) | **Botryosphaeria dieback (239)**  *Botryosphaeria dothidea* (26)  *Diplodia mutila* (98)  *Diplodia seriata* (115)  **Cytospora canker (25)**  *Cytospora viticola* (18)  *Cytospora* sp. (7)  **Esca (47)**  *Cadophora* sp. (3)  *Phaeomoniella chlamydospora* (44)  **Eutypa dieback (14)**  *Cryptovalsa ampelina* (14)  **Phomopsis dieback (1)**  *Diaporthe ampelina* (1) | **Botryosphaeria dieback (288)**  *Diplodia seriata* (288)  **Cytospora canker (190)**  *Cytospora viticola* (190)  **Esca (2)**  *Phaeomoniella chlamydospora* (2)  **Phomopsis dieback (13)**  *Diaporthe* *ampelina* (13) |
| **Esquive** | **Botryosphaeria dieback (107)**  *Botryosphaeria dothidea* (16)  *Diplodia mutila* (62)  *Diplodia seriata* (29)  **Cytospora canker (50)**  *Cytospora predappioensis* (1)  *Cytospora viticola* (43)  *Cytospora* sp. (6)  **Esca (67)**  *Phaeoacremonium* *minimum* (1)  *Phaeoacremonium* sp. (4)  *Phaeoacremonium* *vitivola* (1)  *Phaeomoniella chlamydospora* (61)  **Eutypa dieback (3)**  *Eutypa lata* (3)  **Phomopsis dieback (4)**  *Diaporthe ampelina* (4) | **Botryosphaeria dieback (165)**  *Botryosphaeria dothidea* (8)  *Diplodia mutila* (125)  *Diplodia seriata* (32)  **Cytospora canker (10)**  *Cytospora viticola* (10)  **Esca (46)**  *Phaeomoniella chlamydospora* (46)  **Eutypa dieback (9)**  *Cryptovalsa ampelina* (9) | **Botryosphaeria dieback (256)**  *Botryosphaeria dothidea* (1)  *Diplodia mutila* (29)  *Diplodia seriata* (226)  **Cytospora canker (192)**  *Cytospora viticola* (192) |
| **Madiran** |  |  |  |
| **Control** | **Botryosphaeria dieback (120)**  *Diplodia seriata* (111)  *Neofusicoccum* sp. (9)  **Esca (44)**  *Cadophora luteo-olivacea* (4)  *Phaeomoniella chlamydospora* (40)  **Eutypa dieback (1)**  *Eutypella vitis* (1)  **Phomopsis dieback (16)**  *Diaporthe ampelina* (2)  *Diaporthe foeniculina* (12)  *Diaporthe* sp. (2) | **Botryosphaeria dieback (131)**  *Diplodia sapinea* (32)  *Diplodia seriata* (99)  **Esca (20)**  *Cadophora* sp. (1)  *Phaeomoniella chlamydospora* (19)  **Phomopsis dieback (17)**  *Diaporthe ampelina* (10)  *Diaporthe foeniculina* (5)  *Diaporthe rudis* (1)  *Diaporthe* sp. (1) | **Botryosphaeria dieback (431)**  *Diplodia seriata* (233)  *Neofusicoccum parvum* (198) |
| **Vintec** | **Botryosphaeria dieback (97)**  *Diplodia seriata* (66)  *Neofusicoccum* sp. (31)  **Esca (27)**  *Cadophora luteo-olivacea* (1)  *Phaeomoniella chlamydospora* (26)  **Phomopsis dieback (54)**  *Diaporthe ampelina* (2)  *Diaporthe foeniculina* (21)  *Diaporthe* sp. (21)  *Diaporthe* *neoviticola* (9)  *Diaporthe* *viticola* (1) | **Botryosphaeria dieback (164)**  *Botryosphaeria obtusa* (1)  *Diplodia sapinea* (29)  *Diplodia seriata* (124)  *Neofusicoccum parvum* (10)  **Cytospora canker (1)**  Cytospora sp. (1)  **Esca (17)**  *Cadophora* sp. (3)  *Phaeomoniella chlamydospora* (14)  **Phomopsis dieback (35)**  *Diaporthe ampelina* (28)  *Diaporthe corticola* (1)  *Diaporthe foeniculina* (1)  *Diaporthe* sp. (5)  **Eutypa dieback (3)**  *Eutypella vitis* (3) | **Botryosphaeria dieback (314)**  *Diplodia seriata* (222)  *Neofusicoccum* sp. (92)  **Phomopsis dieback (9)**  *Diaporthe neoviticola* (9) |
| **Tessior** | **Botryosphaeria dieback (94)**  *Neofusicoccum* sp. (13)  *Diplodia seriata* (81)  **Esca (19)**  *Cadophora luteo-olivacea* (1)  *Phaeomoniella chlamydospora* (18)  **Eutypa dieback (5)**  *Eutypella vitis* (5)  **Phomopsis dieback (22)**  *Diaporthe foeniculina* (19)  *Diaporthe* sp. (3) | **Botryosphaeria dieback (169)**  *Diplodia sapinea* (26)  *Diplodia seriata* (100)  *Neofusicoccum parvum* (43)  **Esca (21)**  *Cadophora* sp. (4)  *Phaeomoniella chlamydospora* (17)  **Eutypa dieback (5)**  *Cryptovalsa ampelina* (5)  **Phomopsis dieback (74)**  *Diaporthe ampelina* (40)  *Diaporthe foeniculina* (18)  *Diaporthe melonis* (6)  *Diaporthe phaseolorum* (1)  *Diaporthe rudis* (10)  *Diaporthe* sp. (9)  **Eutypa dieback (3)**  *Eutypella vitis* (3) | **Botryosphaeria dieback (346)**  *Diplodia seriata* (346)  **Phomopsis dieback (5)**  *Diaporthe ampelina* (5) |
| **Bloccade** | **Botryosphaeria dieback (163)**  *Botryosphaeria dothidea* (1)  *Diplodia seriata* (121)  *Neofusicoccum* sp. (41)  **Esca (42)**  *Cadophora luteo-olivacea* (5)  *Phaeomoniella chlamydospora* (37)  **Eutypa dieback (1)**  *Eutypa lata* (1)  **Phomopsis dieback (30)**  *Diaporthe ampelina* (19)  *Diaporthe foeniculina* (5)  *Diaporthe* *rudis* (4)  *Diaporthe* *rumicicola* (2) | **Botryosphaeria dieback (209)**  *Diplodia sapinea* (70)  *Diplodia seriata* (93)  *Neofusicoccum* *parvum* (46)  **Cytospora canker (5)**  *Cytospora* sp. (5)  **Esca (53)**  *Cadophora* sp. (14)  *Phaeomoniella chlamydospora* (39)  **Eutypa dieback (1)**  *Eutypella scoparia* (1)  **Phomopsis dieback (36)**  *Diaporthe ampelina* (21)  *Diaporthe foeniculina* (1)  *Diaporthe* *rudis* (1)  *Diaporthe* *nigra* (13) | **Botryosphaeria dieback (290)**  *Neofusicoccum* sp. (290)  **Phomopsis dieback (2)**  *Diaporthe neoviticola* (2) |
| **Blindar** | **Botryosphaeria dieback (219)**  *Diplodia seriata* (219)  **Esca (23)**  *Cadophora sp.* (2)  *Phaeomoniella chlamydospora* (21)  **Eutypa dieback (4)**  *Eutypella vitis* (4)  **Phomopsis dieback (62)**  *Diaporthe ampelina* (18)  *Diaporthe foeniculina* (19)  *Diaporthe* *melonis* (3)  *Diaporthe* *rudis* (3)  *Diaporthe* sp. (12)  *Diaporthe* *viticola* (7) | **Botryosphaeria dieback (372)**  *Botryosphaeria dothidea* (1)  *Diplodia sapinea* (84)  *Diplodia seriata* (228)  *Neofusicoccum parvum* (59)  **Esca (19)**  *Cadophora* sp. (2)  *Phaeomoniella chlamydospora* (17)  **Phomopsis dieback (67)**  *Diaporthe ampelina* (53)  *Diaporthe foeniculina* (12)  *Diaporthe nigra* (2) | **Botryosphaeria dieback (194)**  *Diplodia seriata* (42)  *Neofusicoccum* sp. (152) |
| **Esquive** | **Botryosphaeria dieback (124)**  *Diplodia seriata* (124)  **Esca (38)**  *Cadophora sp.* (2)  *Phaeomoniella chlamydospora* (36)  **Eutypa dieback (1)**  *Cryptovalsa ampelina* (1)  **Phomopsis dieback (36)**  *Diaporthe ampelina* (11)  *Diaporthe foeniculina* (6)  *Diaporthe* *nigra* (3)  *Diaporthe* *rudis* (3)  *Diaporthe* sp. (12)  *Diaporthe* *viticola* (1) | **Botryosphaeria dieback (131)**  *Botryosphaeria dothidea* (2)  *Diplodia sapinea* (23)  *Diplodia seriata* (60)  *Neofusicoccum parvum* (46)  **Eutypa dieback (3)**  *Eutypa lata* (3)  **Esca (22)**  *Cadophora sp.* (9)  *Phaeomoniella chlamydospora* (13)  **Phomopsis dieback (20)**  *Diaporthe ampelina* (9)  *Diaporthe foeniculina* (1)  *Diaporthe* *melonis* (1)  *Diaporthe* *nigra* (4)  *Diaporthe* *rudis* (2)  *Diaporthe* sp. (3) | **Botryosphaeria dieback (225)**  *Diplodia seriata* (225) |

^a^Number of isolates.
